# Reduced baseline male fertility and increased sensitivity to hypoxic stress in high-elevation deer mice

**DOI:** 10.64898/2026.10.16.759662

**Authors:** Kelsie E. Hunnicutt, Emma S. Glenn, Joe C. Perry, Madison Q. Tatro, Lacey K. Conrad, Makenna Y. Juergens, Kathryn Wilsterman, Benjamin M. Skinner, Erica L. Larson, Jonathan P. Velotta

**Affiliations:** Department of Biological Sciences, University of Denver, Denver, CO 80210, USA; Department of Biology, Stanford University, Stanford, CA 94305, USA; School of Life Sciences, University of Essex, Colchester CO4 3SQ, UK; Department of Biology, Colorado State University, Fort Collins, CO 80523, USA

**Author notes:** **Corresponding Author:** Kelsie Hunnicutt, Department of Biology, Stanford University, Stanford, CA 94305.

**Keywords:** elevation, male fertility, spermatogenesis, *Peromyscus*, gene expression

## Abstract

Extreme environments like high elevation impose chronic challenges to endothermic animals who must sustain metabolic demand despite reduced oxygen availability. Male reproduction is a tightly regulated physiological system, and hypoxia exposure has negative but transient effects on male reproductive output in diverse mammal species not adapted to high-elevation. Little is known about whether high elevation populations adapt to the detrimental effects of hypoxia on male reproductive physiology. Here, we use high- and low-elevation populations of deer mice (*Peromyscus maniculatus*) and a lowland congener, *P. leucopus,* to test how male reproductive phenotypes have diverged across populations and in response to hypoxia exposure across acute, intermediate, and chronic timescales. We complement these phenotypic assays with bulk RNA sequencing of testes to assess the testis-specific response to hypoxia. Contrary to expectations, hypoxia exposure had a negligible effect on low-elevation strains, while highland mice had both lower baseline fertility and heightened sensitivity to hypoxic stress including complete absence of mature sperm. Gene expression patterns indicate that hypoxia exposure dramatically reorganizes testes architecture in highland mice, consistent with testicular regression similar to that observed in seasonally breeding mammals. We hypothesize that this increased sensitivity of male reproductive physiology in highlanders reflects a life history trade-off, allowing energy reallocation between reproduction, survival, and thermogenesis in response to stress in extreme environments.

## Introduction

The extremely low oxygen availability of high-elevation environments applies intense selective pressure on resident endothermic animals who must continuously supply oxygen to respiring tissues. The physiological constraints imparted by low oxygen conditions at high elevations can lead to physiological dysfunction and decreased fitness (Swenson & Bartsch, 2016). For example, sustained red blood cell production in response to hypoxia leads to elevated blood viscosity, which can burden the cardiovascular system (Storz, Scott, et al., 2010). As a result, high-elevation resident populations have evolved suites of adaptations to mitigate the detrimental effects of chronic hypoxia including more efficient delivery of oxygen to tissues through shifts in hemoglobin-oxygen binding (Hanaoka et al., 2012; Peng et al., 2017; Storz, Runck, et al., 2010; Velotta et al., 2025) and resistance to cardiovascular dysfunction (West et al., 2021). Other studies have focused on adaptations of female reproduction including changes in placental anatomy (Arenas et al., 2026; Wilsterman et al., 2023) that protect newborn offspring from the restrictive effects of hypoxia on fetal growth. In sum, high-elevation environments provide an excellent testing ground for studies of the mechanisms of adaptation.

While a large body of work has uncovered adaptations of oxygen transport systems at high elevation and of female reproduction, very few studies have investigated whether male reproductive physiology evolves in high-elevation animals (*e.g.,* in Tibetan sheep; (H.-H. Wang et al., 2026)), despite the sensitivity of spermatogenesis to oxygen availability. Spermatogenesis begins with mitotically dividing spermatogonia of two main types, self-renewing spermatogonial stem cells and progenitor cells that differentiate into spermatocytes, enter meiosis, and produce haploid spermatids. As germ cells progress through spermatogenesis, their requirements for available oxygen shift. Spermatogonial stem cells occupy the lowest oxygen-containing compartments of the testis (Lord, 2025; Lord & Nixon, 2020), primarily use glycolysis for self-renewal (Helsel et al., 2017; Kanatsu-Shinohara et al., 2016), and rely on expression of the hypoxia-inducible transcription factors *Epas1* and *Hif1a* to maintain regenerative capacity (Bernstein et al., 2023; Morimoto et al., 2021). In contrast, progenitor spermatogonia and spermatocytes require more available oxygen to fuel oxidative phosphorylation, and the highest levels of oxygen availability are needed for the final stages of sperm maturation and eventually fertilization (Bajpai et al., 1998; Lord, 2025).

Given the dependency of developing germ cells on oxygen availability, exposure to high-elevation (*i.e.,* hypobaric) hypoxia, either naturally or simulated in lab environments, predictably impairs sperm production in a variety of mammalian systems. This includes human visitors and migrants to high-elevation environments (Gonzales, 2007; J. He et al., 2015; Verratti et al., 2016, 2022) as well as laboratory colonies of rodents (S. Li & Yang, 2022; Liao et al., 2010; Tao et al., 2022) and rhesus monkeys (Saxena, 1995). Negative effects on male reproductive physiology can arise after as few as seven days of exposure to hypobaric hypoxia (Tao et al., 2022) and include decreased quantity of sperm produced, impaired sperm motility and viability, and reduced fertilization potential (Lord, 2025; Reyes et al., 2012; Zhong et al., 2024). Paternal hypoxia exposure can even impact male offspring reproductive physiology in house mice (S. Li & Yang, 2022). However, given either sufficient time to acclimate to hypoxia exposure or a return to normal oxygen conditions, fertility of non-adapted organisms often recovers to pre-exposure levels, although full recovery can take at least 6-9 weeks in rodents (Bai et al., 2018; S. Li & Yang, 2022; Z. Li et al., 2021), and some studies in humans suggest that full recovery may take between several months and two years (Okumura et al., 2003; Verratti et al., 2022).

It is unclear whether and how populations that live permanently at high elevation have evolved to respond to hypoxic stressors on male reproduction (Swenson & Bartsch, 2016; Yu et al., 2025). Given the detrimental effects of hypobaric hypoxia on sperm production in lowland resident populations, we might expect that long-term high-elevation populations would experience selective pressures for physiological adaptations that ensure proper supply of oxygen to developing sperm and buffer male reproduction against hypoxic stress. Deer mice (*Peromyscus maniculatus*) are a well-suited model system for investigating how male reproductive physiology can adapt to high-elevation environments. *P. maniculatus* has the broadest elevational distribution of any North American mammal (Hock, 1964), and the genetic mechanisms underlying adaptations to high elevation are well characterized for many biological processes (Storz et al., 2019). Additionally, deer mice have a promiscuous mating system in which females mate with multiple males (Fisher et al., 2014; McLean et al., 2022), which should provide strong selective pressure to maintain male reproductive physiology (as in Weber & Fisher, 2023). High-elevation populations of *P. maniculatus* also maintain large effective population sizes (Schweizer et al., 2021), but direct measurements of whether and how highland males sustain sperm production in low-oxygen conditions are still lacking.

Here, we conducted hypoxia acclimation experiments spanning acute, intermediate, and chronic time scales using lab strains derived from low- and high-elevation populations of *P. maniculatus* and a closely related low-elevation species, *P. leucopus*. We tested the hypothesis that male reproduction in high-elevation deer mice should be buffered against the effects of hypoxia exposure while low-elevation mice should show hypoxia-induced defects in male reproductive physiology. Contrary to our prediction, highland males had reduced baseline fertility and were more sensitive to hypoxia exposure than lowland males, culminating in a complete absence of mature sperm in a subset of individuals in as few as two weeks of hypoxia exposure. We found patterns of gene expression in the testes consistent with a decrease in late spermatogenic cell types and increase in apoptosis in sterile males consistent with a dramatic reorganization of testis architecture and a cessation of spermatogenesis. We identify candidate genes implicated in testosterone production and tissue remodeling whose expression is associated with decreased sperm production in highland mice in response to hypoxia exposure in the testes. Our study provides the first direct comparison of how low- and high-elevation populations differently mitigate the system-wide pressures of low oxygen availability in the male reproductive system. We propose that highland deer mice have evolved an increased sensitivity to environmental stress as a balancing mechanism to allow males to effectively manage energy expenditures between survival and reproduction.

## Methods

### Animal strains and hypobaric hypoxia experimental design

For our initial experiment (hereafter the focal experiment), we used lab colonies of high- and low-elevation populations of *P. maniculatus* originally derived from the summit of Mount Blue Sky, CO (MBS; elevation 4,348 m; 39.5883° N, −105.6438° W) and Kearney, Nebraska (KN; elevation 655.9 m; 40.7864° N, –99.0836° W; Figure 1). Colony founders were collected in 2018 and 2019. We used fourth (KN) and fifth (MBS) generation descendants of founders who were born at Colorado State University (CSU; 1,525 m above sea level) before being transferred to the University of Denver (∼1,640 m above sea level) where they acclimated for > 3 months prior to sampling. For an extended comparison of high- and low-elevation strains, we included additional deer mouse lines from the *Peromyscus* Genetic Stock Center for a low-elevation strain of *P. maniculatus* derived from Ann Arbor, Michigan (BW), a high-elevation strain of *P. maniculatus* derived from the White Mountains of California (SM2), and a closely-related low-elevation strain of the sister species *P. leucopus* originally trapped in Linville Falls, NC (LL; Figure 1; Lucius et al., 2021). Stock center mice were born at low-elevation in South Carolina, transferred to the University of Denver and acclimated to lab conditions for approximately four weeks before experimentation. All mice in the focal and extended comparisons were provided with food and water *ad libitum* and maintained on 12L:12D daylength at room temperature (∼25°C). All animal use for the hypoxia exposure trials was carried out at the University of Denver under IACUC protocols #2213708.

**Figure 1:**
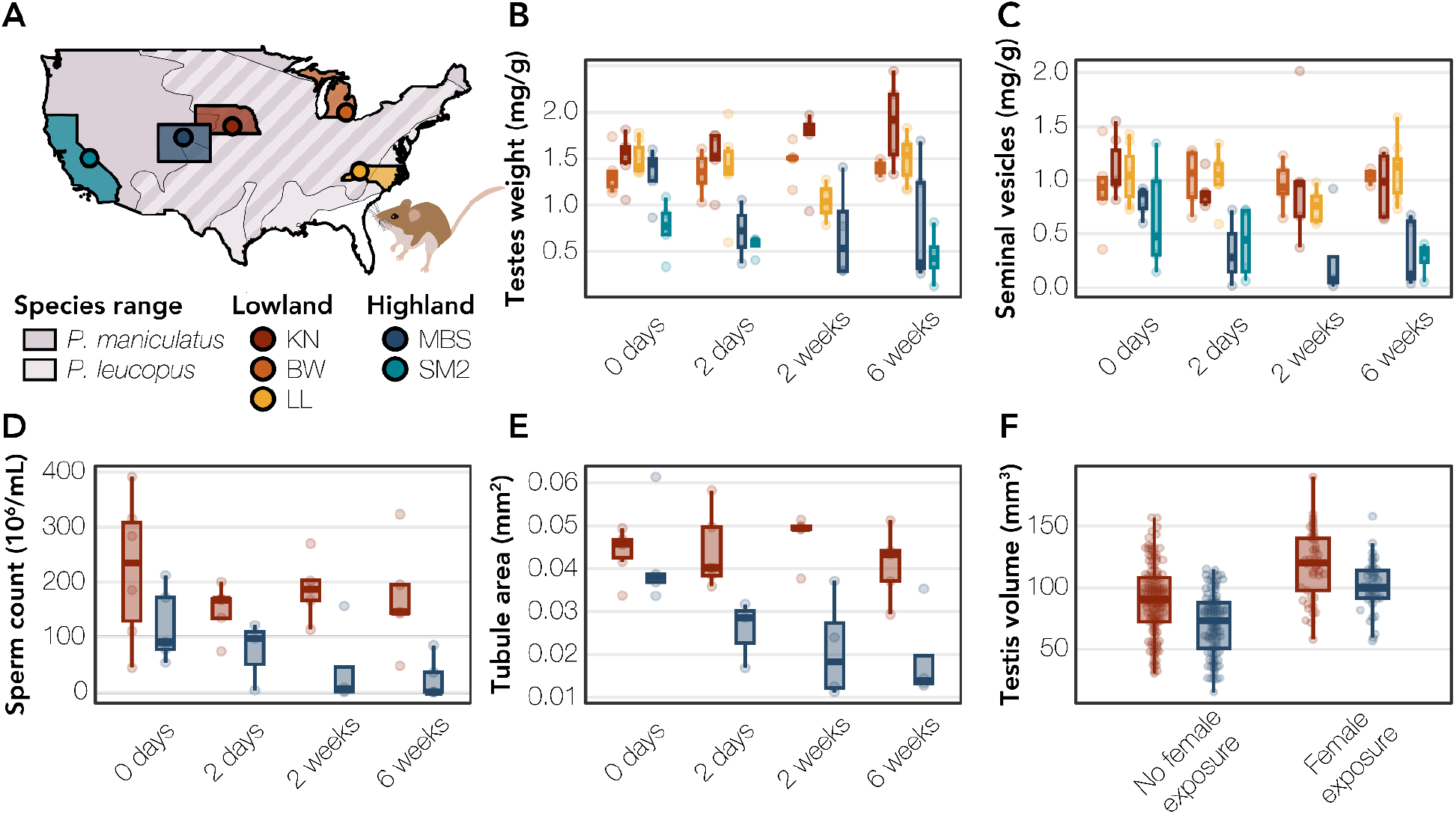
Male reproductive physiology of high- and low-elevation populations of *P. maniculatus* and low-elevation *P. leucopus.* (A) Collection locations and species ranges of lab-derived strains used in the study. Species ranges are based on IUCN Red List estimates (2016, 2017). (B) Relative paired testes weight normalized by body mass across strains and hypoxia exposure durations (control, two days, two weeks, or six weeks). For all boxplots, the median is indicated with whiskers extending to either the largest or smallest value or no further than 1.5 times the interquartile range. (C) Relative paired seminal vesicle weights normalized by body mass. (D) Normalized sperm counts (in 10^6^ million sperm/mL). (E) Average seminiferous tubule area. (F) Average testis volume for deer mice in the CSU colony. Mice were placed in two female exposure categories - unexposed to females and housed only with same-sex siblings and those who were exposed to females and successfully sired offspring after six weeks. Colors indicate strain with cooler colors indicating strains derived from highland populations (MBS = dark blue; SM2 = light blue), and warmer colors indicate mouse strains derived from low-elevation populations (KN = red; BW = orange; LL = yellow).

To disentangle the effects of acute, intermediate, and long-term exposure to hypoxia on male reproductive phenotypes, we conducted two experimental trials in which we randomly assigned reproductively mature males from each strain to treatment groups corresponding to 0 days, 2 days, ∼2-3 weeks, and 6 weeks of hypobaric hypoxia exposure (Table S1) mimicking 4,300 m above sea level (pO2 = 61 kPa) using custom hypobaric hypoxia chambers described elsewhere (Lui et al., 2015). As deer mice from our MBS and KN colonies are more recently derived from the wild (generations F5 and F4 respectively), we assigned siblings from the same families across acclimation treatments to account for family-specific effects (Table S2). The first trial consisted only of KN and MBS mice, while the second trial included the strains in our extended comparison (SM2, LL, and BW). In this second trial, we excluded SM2 mice from the 2-3 week treatment due to low availability of this strain from the stock center. Deer mice varied in age across strains due to limited availability. At the time of dissection, highland mice were on average 351 +/- 8 dpp (MBS; mean +/- 1 SEM) and 582 +/- 36 dpp (SM2) at the time of the experiment, while lowland mice were on average 365 +/- 6 dpp (KN), 404 +/- 19 dpp (BW), and 224 +/- 10 dpp (LL). Because the variation in age was primarily across strains rather than within strains, we were unable to account for the impact of mouse age on reproductive phenotype. To reduce the effects of dominance interactions (Snyder, 1967), we housed mature males from all strains individually for at least two weeks prior to either sample collection or hypoxia exposure. At the time of sampling, we euthanized males using isoflurane overdose and decapitation and immediately processed individuals for male reproductive phenotyping and tissue collection. We used between 3-6 mice per strain per hypoxia exposure treatment (Table S1).

### Male reproductive phenotyping

For all strains, we collected morphometric data (length of right ear, left foot, body, and tail as well as body mass) and dissected out and weighed the seminal vesicles, cauda epididymis, right and left testes, white gonadal fat pads, and the brown adipose pad between the shoulder blades. From this sampling effort, we calculated male fertility metrics including paired testes weight and paired seminal vesicle weight and standardized paired testes weights and seminal vesicle weights by body mass. We fixed one testis from each mouse using Bouin’s fixative (Poly Scientific R&D Corp.) and subsequently washed each testis 24 hours later in a series of ethanol dilutions (25% EtOH twice, 50% EtOH, and 70% EtOH). We stored fixed testes at 4°C until shipment to the Research Histology Service at the University of Colorado (CU) at Anschutz for processing. Briefly, each testis was bisected across the short axis and embedded in paraffin wax. Two slides of three 5µm paraffin-embedded samples each were produced, discarding 10µM between sections. Samples were stained for visualization of seminiferous tubule morphology with Hematoxylin and Eosin (H&E). We imaged each testis using a ZEISS Primostar 3 microscope at 100-200x. For each individual, we measured a minimum of ten seminiferous tubules and recorded the diameter of the long axis, diameter of the short axis, tubule perimeter, and tubule area using the Labscope software (ZEISS) with the line and spline tools (Table S3).

For mice in our focal comparison, we dissected and flash-froze one testis for subsequent RNA sequencing and conducted additional reproductive phenotyping including sperm counts and morphometrics. We estimated sperm count from the caudal epididymides following Good et al. (2008) by dicing the tissue in 1ml of Dulbecco’s PBS and incubating at 37°C for 10 minutes. We counted intact sperm (both head and tail present) in a heat shocked 5uL suspension across a fixed area (5 rows) of a Makler counting chamber (IVF Store). After sperm counts, we prepared the remainder of the extracted caudal sperm for sperm morphometrics following Mandawala et al. (2026). We diluted the caudal extraction in 3mL PBS, centrifuged for 10 minutes at 1500 g, washed the cell pellet twice with 2mL PBS, and centrifuged the resuspended pellet at 1500 g for 5 minutes. We removed the supernatant, thoroughly resuspended the pellet, and fixed sperm in 500 uL cold 3:1 Methanol: Acetic acid. We stored fixed sperm at -20C and shipped samples to the University of Essex for subsequent analysis.

To assess differences in male reproductive phenotypes across strains, we ran linear mixed effects models using the lme4 v. 2.0-6 and lmerTest v. 3.2-1 package in R v. 4.5.0. For relative testes weight and relative seminal vesicle weight, we ran three models each: (1) a full model with mouse strain, acclimation treatment, and their interactions as the main effects, (2) an elevation model, with elevation of strain origin (high vs low), acclimation treatment, and their interactions as the main effects, and (3) a focal model, with mouse strain (only MBS and KN), acclimation treatment, and their interactions as the main effects (Tables S4 - S6). We included mouse family nested within strain as a random effect to account for family-specific differences in reproductive phenotypes. We measured normalized sperm counts and average semeniferous tubule area only in our focal comparison, and so we analyzed these traits with only the focal model.

### Sperm morphometrics

From each sample, we dropped 4µl of 3:1 Methanol: Acetic acid fixed sperm on a slide and stained sperm with 16μl VectorShield Antifade Mounting Medium with DAPI (Vector Labs) under a 22x50mm cover slip. We imaged slides using a 20x20 field area on a Nikon Ti2 widefield fluorescence microscope equipped with a Kinetix A22M723026 camera (Teledyne) using a Plan Apo λ 100x oil objective lens with 1.5X tube lens active. We analyzed sperm morphology from images using Nuclear Morphology Analysis v. 2.5.0 (Skinner, 2022; Skinner et al., 2019) and characterized sperm into broad morphometric groups as described in Mandawala et al. (2026). Briefly, we clustered cells by shared abnormalities in regions of the sperm head to yield three groups: normal, intermediate, and highly abnormal sperm. We identified these clusters using nonunimodal regions clustered via K-means and separated clusters into feature groups using the hamming distance between representative codes. We then inspected these groups and used NMA to characterise them. We visualized overall sperm shape distributions using t-SNE in the Rtsne package on morphological profiles that were significantly different (p <0.05) in R v. 4.40 (Figure S1). We performed permutational multivariate analysis of variance (PERMANOVA; using the *adonis2* function in the vegan package) on the differences between populations on an individual scale with proportion of cells in each category per individual. To verify the assumption of homogeneous multivariate dispersion, we analyzed distances among samples using betadisper and followed this with analysis of variance.To identify which morphological categories contributed to population differences, we fit separate linear models for the percentages of normal, intermediate, and abnormal nuclei, with population and hypoxia treatment duration included as explanatory variables. In addition, we tested populations for pairwise differences for each morphological category using two-sided Wilcoxon rank-sum tests. We assessed population differences across the complete multivariate morphological profile dataset using multivariate analysis of variance (MANOVA) with Pillai’s trace statistic. We subsequently analyzed individual profile variables using univariate linear models with population as the explanatory factor, and corrected p-values for multiple testing using both Benjamini-Hochberg false discovery rate (FDR) and Bonferroni procedures. We quantified variability using interquartile ranges (IQRs) calculated separately for each population. To identify which morphological categories contributed to population differences, we fit separate linear models for the percentages of normal, intermediate and abnormal nuclei, with population and hypoxia treatment duration included as explanatory variables. Population-specific differences within each morphological category were additionally assessed using two-sided Wilcoxon rank-sum tests. We evaluated differences in variance between populations for each morphological category using Fligner-Killeen tests and summarized variability using standard deviations and interquartile ranges.

### Repeated estimation of baseline male fertility

To confirm the baseline fertility results from our initial experiment, we analyzed a large, opportunistically collected phenotypic data set that included testis size measurements from wild-derived lab-reared (F1-F3 generation) individuals at Colorado State University (CSU). These animals were derived from a different set of wild parents captured in Kearney, Nebraska and Mount Blue Sky, Colorado from 2023 - 2025. Deer mice in this colony were housed post-weaning with same-sex conspecifics on 14L:10D daylength at room temperature (∼23°C) and euthanized as above under ambient, normobaric conditions in the lab. We collected testes measurements from males from three categories: (1) males housed only with other males (*i.e.*, same-sex conspecifics; n = 143 KN and n = 101 MBS males), and (2) males paired with a female who successfully sired offspring after 6 weeks (n = 45 KN and n = 40 MBS). Male mice housed only with other males were on average 116 +/- 3 dpp (KN) and 119 +/- 3 dpp (MBS) at time of euthanasia, while males housed with females were 225 +/- 9 dpp (KN) and 217 +/- 8 dpp (MBS). We collected testis size measurements using electronic calipers to measure the length and width of each testis. We calculated testes volume using the equation for the volume of an ellipsoid: ⁴⁄₃Π × (½ length) × (½ width) x (½ height), where width and height were given the same value and averaged the two volumes within each individual. Animal work completed at Colorado State University was covered under IACUC-approved protocols 2895 and 6657.

### Bulk RNA sequencing of testes

For mice in our initial focal comparison, we extracted RNA from ∼25-50 mg of testis using Qiagen RNEasy Mini kits (Qiagen) with Trizol reagent (Invitrogen). We homogenized tissues in 1mL Trizol in a Powerlyzer or Tissuelyzer (Qiagen) with ceramic 2.8 mm Beads in Phasemaker tubes (Invitrogen). We sequenced RNA from samples through Novogene, requiring a RIN > 7 which was measured by Novogene on a 5400 Fragment Analyzer System (Table S7). Novogene prepared sequencing libraries using KAPA HyperPrep kits (Roche) and sequenced each library on an Illumina NovaSeq X Plus with 150 bp paired-end sequencing. This resulted in 3-5 samples per strain and hypoxia exposure treatment (Table S7).

### Gene expression pre-processing

We generated RNA sequencing data of whole testes resulting in an average of ∼28.0 million raw read pairs per individual (Table S7). We trimmed raw reads of adapters and low quality bases and reads using Trimmomatic v.0.39 (Bolger et al., 2014) and removed low quality bases from the first and last 5 bp of each read and bases with an average Phred score of <15 across a 4 bp sliding window. We only retained reads of at least 36 bp. We mapped trimmed reads with STAR v. 2.7.11b (Dobin et al., 2013) to the closest available reference genome (*P. maniculatus* representing strain BW (GCF_049852395.1; Gozashti et al., 2025) with the associated NCBI RefSeq annotation and quantified read counts per gene using default settings. On average, 78.53% (KN) and 78.72% (MBS) of read pairs mapped uniquely per sample (Table S7). We imported read counts into R v. 4.5.0 for differential expression and gene network analysis. We filtered out lowly expressed genes using the filterByExpr command from edgeR v. 4.6.3 (Y. Chen et al., 2016) requiring a gene to have a minimum of 10 counts normalized by library size independently within each strain. After filtering, we retained 20,588 genes in the expressed gene set.

### Differential expression

To characterize broad patterns of gene expression across strains and treatments, we generated multidimensional scaling (MDS) plots using the plotMDS function in limma v. 3.64.3 (Ritchie et al., 2015) using the top 1,000 genes driving expression differences between treatments. We used edgeR to normalize read counts across libraries and estimated differential expression in response to strain, hypoxia exposure, and the interaction between strain and hypoxia exposure. We constructed a design matrix using a full factorial model (strain × hypoxia exposure). We fit the dataset with genewise negative binomial generalized linear models with quasi-likelihood dispersion estimation (Lund et al., 2012). For each factor (strain, hypoxia exposure, and their interaction), we performed a quasi-likelihood F-test with glmQLFTest. We modeled hypoxia exposure as a time course, with exposure duration as an unordered categorical factor with four levels: 0 days (the reference), 2 days, 2 weeks, and 6 weeks. We coded strain as a two-level factor with KN as the reference. We considered genes differentially expressed if their absolute log2 fold-change was greater than 1 and if the Benjamini-Hochberg corrected-false discovery rate (FDR) was below 0.05 (Benjamini & Hochberg, 1995). We assessed differentially expressed gene sets for each fixed effect for enrichment in different Gene Ontology (GO) terms using gProfiler2 v.0.2.4 (Kolberg et al. 2020) in R. We only included genes expressed above our expression threshold as the background in our GO analysis. We retained Molecular Function (MF), Cellular Component (CC), and Biological Process (BP) GO terms with an FDR below 0.05 and ran gProfiler2 both with and without the highlight option, a two-stage algorithm for reducing resulting GO terms by grouping significant terms into sub-ontologies and then identifying the gene sets that give rise to other significant functions (Kolberg et al. 2020). Finally, we looked for overlap between differentially expressed genes and a gene set with elevated lineage-specific evolution in wild MBS individuals based on the population branch statistic from a previous study that used whole exome data (*i.e.,* PBS outliers; Schweizer et al., 2021; Wilsterman et al., 2023).

### Cell type deconvolution and apoptosis gene expression

To assess testis samples for potential shifts in cellular composition, we used an *in silico* bulk RNAseq deconvolution approach as implemented in MuSiC v. 1.0.0 (X. Wang et al., 2019) using a testis single cell RNAseq reference atlas from adult house mice (GEO accession: GSE112393; Green et al., 2018). We used the ST1-ST8 samples from the atlas and their per-cell cell type annotations which included the following cell types: innate lymphocytes, endothelial cells, myoid cells, Leydig cells, Sertoli cells, an unresolved (“unknown”) population, spermatogonia, spermatocytes, round spermatids, and elongating spermatids. We then estimated cell type proportions from our bulk RNAseq data using the *music_prop* function which produced two estimates of cell-type proportions per sample: a MuSiC-weighted estimate (Est.prop.weighted) and a non-negative least squares (NNLS) estimate using all genes (Est.prop.allgene). To test for differences in cell type proportions, we used beta regression in the betareg v. 3.2-6 package to model the effects of strain, hypoxia exposure duration, and their interaction for the following cell types: Sertoli cells, spermatogonia, spermatocytes, and elongating spermatids. Prior to regression, we adjusted cell type proportions with the Smithson-Verkuilen transformation to account for individuals with exact zero proportions (Smithson & Verkuilen, 2006). We rescaled sperm count (divided by 1,000) prior to fitting, and adjusted p-values with false discovery rate correction (Benjamini & Hochberg, 1995). We used ComplexHeatmap v.2.24.1 (Gu et al., 2016) to visualize differences across all cell type proportions across samples (Supplemental Figure S3).

Because of potential shifts in cell composition across samples, we next tested for shifts in the expression of apoptosis genes in response to hypoxia exposure. For this analysis, we were interested in strain-specific shifts in genes belonging to the parent apoptosis GO term GO:0006915 (n = 2,082 genes of which n = 1,632 were expressed in our dataset) for each hypoxia exposure group relative to the strain-specific control mice (0 days). We created a separate DGE object with samples assigned to its strain x hypoxia exposure group. For each strain, we defined contrasts comparing the hypoxia exposure groups to the strain-specific control group resulting in six contrasts total. Using edgeR, we estimated dispersions using estimateDisp, fit the model with glmQLFit, and estimated significance with quasi-likelihood F-tests with glmQLFtest. We used this DGE object to test whether the apoptosis gene set as a whole exhibited a coordinated shift in expression across hypoxia exposure groups using the competitive gene set test in *cameraPR* from the limma v. 3.64.3 package. We corrected p-values for multiple testing using the false discovery rate (Benjamini & Hochberg, 1995). For each hypoxia exposure group, we visualized the log fold-change of apoptosis genes in each group relative to its strain-specific control.

### Testes gene co-expression networks

We performed weighted gene correlation network analysis (WGCNA; Langfelder & Horvath, 2008) in R combining MBS and KN individuals (n = 39 individuals). This approach clusters genes with correlated expression patterns into ‘modules’ and summarizes the expression of each module by calculating module ‘eigengene’ scores which represents the first principal component of the standardized expression matrix for genes within a module. We detected modules with the *blockwiseModules function* with the networkType set to “signed” and used the dynamic tree-cutting method (Langfelder & Horvath, 2008). We chose a soft thresholding power of β = 8 to achieve an approximately scale-free topology by assessing the point at which the scale-free topology model fit began to plateau. We ran WGCNA using normalized cpm values for expressed genes and identified modules whose eigengenes were correlated with normalized sperm count as well as modules significantly associated with strain, acclimation treatment, or interaction effects using lmer in R. We performed functional enrichment analysis of GO terms of genes within modules with either a significant correlation to normalized sperm counts or with a significant effect of strain, acclimation, or interaction using gProfiler2 as described above. Background genes sets for module GO enrichment were the genes that passed the WGCNA expression filters for inclusion within the network. For each module, we identified “hub” genes, or genes whose expression within a module is highly correlated with other genes within the module (eigengene-based connectivity or kME). We classified genes as hub genes if their kME was in the top 5% for a given module and annotated hub genes that were also present in the previous dataset of genome-wide outliers for sequence differentiation between highlanders and lowlanders.

## Results

### Baseline male reproductive physiology differences between lowland and highland mice

To assess the effects of acute and chronic hypobaric hypoxia exposure on male reproductive physiology, we examined differences between highland and lowland deer mice in a suite of traits including relative paired testes weight, normalized sperm count, relative seminal vesicle weight, seminiferous tubule area, and mature sperm morphology (Figures 1B-1E and 2). All male reproductive phenotypes were significantly positively correlated (ρ: 0.59 - 0.99; Figure S2). For the extended comparisons of all five strains (low: KN, BW, LL; high: MBS and SM2), we used two linear mixed effects models, one which tested all five strains separately (full model) and a second which tested for the effects of elevation with strain as a nested effect of elevation (elevation model). For the extended comparison, we could only assess differences in relative testes weight and relative seminal vesicle weight. For the focal comparison (focal model: MBS vs KN), we directly contrasted these two strains for normalized sperm count, average semeniferous tubule area, relative testes weight, and relative seminal vesicle weight. The full results of the linear mixed effects models are included in Table S4 - S6.

We first tested whether there were differences in baseline reproductive physiology across strains and found trait-specific evidence that baseline reproductive output was reduced in high-elevation strains. SM2 individuals, but not the MBS individuals, had significantly lower relative testes weights compared to lowlanders in the full model (β = −0.58, p = 0.004). In the elevation model, which considers strains derived from similar elevations together, high-elevation mice had reduced relative seminal vesicle weight compared to low-elevation mice (β = −0.30, p = 0.01). In the focal comparison, the MBS strain did not differ significantly from the lowland KN strain for baseline relative testes weight, seminal vesicle weight, or seminiferous tubule area, but did have marginally lower normalized sperm count (β = -93.7, p = 0.07). In contrast, neither of the low-elevation strains (KN and LL) showed reduced reproductive output compared to the reference strain (BW) in the full model. In sum, across our first experiment, we found mixed evidence of trait- and strain-specific reductions in baseline fertility in high-elevation deer mice.

Testes mass data collected at a replicate deer mouse colony (Fort Collins, CO) showed a similar pattern to the focal comparison: MBS mice had smaller testis volumes compared to KN mice, and this difference persisted in animals that were exposed to females and successfully sired offspring in breeding pairs (Figure 1F). After accounting for body length (F1, 321 = 10.7, p = 0.001) and body mass (F1, 321 = 29.1, p < 0.001), our full linear model revealed a significant main effect of population (F1, 321 = 46.7, p < 0.001) on average testis volume. Additionally, female exposure (F1, 321 = 61.8, p < 0.001) had a significant effect on testis volume resulting in larger testes volumes in males from both populations relative to those housed only with same-sex siblings (Figure 1F). These data support the finding from our focal and extended comparisons that high-elevation mice have reduced testis size compared to lowlanders at baseline.

### Strain-specific responses of male reproductive physiology to hypoxia exposure

We tested whether any of our hypoxia exposure treatments had an effect on male reproductive physiology across strains and/or elevations and found minimal evidence for a general effect of hypoxia exposure on deer mouse reproductive physiology. In all three models, there was no main effect of hypoxia exposure for any of the three durations for either relative testes weight or seminal vesicle weight (Table S4 - S6). The one exception to this was a significant negative effect of two day hypoxia exposure (β = -85.8, p = 0.05) within the focal model only. This suggests that in general, hypoxia exposure impacts deer mouse reproductive physiology in a strain- and/or elevation-specific way across different intervals of hypoxia exposure.

We next tested for a low-elevation strain-specific response to hypoxia exposure. Against the BW and control exposure, the low-elevation strains showed a minimal response to hypoxia. As discussed above, in the focal comparison, there was a significant negative effect of 2 days of hypoxia exposure on normalized sperm counts although this was not strain specific. However, most of the strain-by-hypoxia exposure interaction terms for relative testes weight and seminal vesicle weight in the full model were non-significant for KN and LL (Table S4), with the exception of a significant decline in relative testes weight at two weeks in only the LL strain (β = -0.60, p = 0.03).

In contrast to the low-elevation strains, the high-elevation strains showed a markedly stronger and more consistent impairment of male reproductive physiology in response to hypoxia exposure. In the elevation model, high-elevation mice exhibited marginal or significant exposure-dependent declines in relative testes weight at two days (β = -0.35, p = 0.08), two weeks (β = -0.44, p = 0.06), and six weeks (β = -0.57, p = 0.003), and a parallel trend in seminal vesicle weight decline, especially at six weeks (β = -0.45, p = 0.01). The MBS strain showed this pattern of hypoxia-induced fertility declines most clearly in the focal model, with significant reductions compared to KN in relative testes weight (two weeks: p = 0.04; six weeks: p = 0.02) and average seminiferous tubule area (two days: p = 0.06; two weeks: p = 0.005; and six weeks: p = 0.02), although relative seminal vesicle weight was non-significant across all three durations (Table S5). Although normalized sperm count showed shared general declines in response to the three hypoxia treatments in the focal comparison, complete azoospermia arose in multiple MBS individuals at two weeks (n = 1/4 mice) and six weeks (n = 3/5 mice). No KN mice in any treatment showed similar reproductive impairment. Taken together, our results indicate that duration of hypoxia exposure, in addition to baseline strain differences, result in impaired reproductive physiology in high-elevation mice and that this reduced reproductive output in highland mice is a derived phenotype relative to the ancestral lowland *P. leucopus* strain and lowland *P. maniculatus* strains.

Finally, we tested whether highland and lowland deer mice exhibit altered spermiogenesis (the final stages of sperm maturation) in response to hypoxia exposure through an assessment of mature sperm morphology. We visualized morphological variation across all measured nuclei using tSNE and used clustering to identify the broad morphological phenotypes. The morphological analysis captured two primary axes of morphological variation. One axis was consistent with a perturbation of spermiogenesis (Figure 2A & 2C) and the other was consistent with subtle variation of nuclear shape between the two populations (Figure 2C). The axis of subtle inter-population morphological difference was identified via a MANOVA (p < 2.2e^-16^) on morphological profiles with Bonferroni correction, finding 276 morphological points with significant differences (P < 0.05). We then visualized the differences in morphospace using tSNE (Fig 2C). The axis consistent with perturbation of spermiogenesis was split into three morphological groupings: normal, containing cells with no signs of morphological difference from the expected nuclear shape; intermediate, which show some subtle signs of morphological abnormality; and abnormal, which show more obvious signs of abnormality. For each individual, we assessed variability in sperm nuclear morphology across the three groups (normal, intermediate and abnormal) and found a significant difference (p=0.017) between variability of the proportion of abnormal sperm nuclei. We found no difference between strains in the proportion of normal sperm (p=0.76) or intermediate sperm (p=0.095), but the KN population had more variable proportions of abnormal sperm (IQR ratio 2.18, SD 1.57). In contrast, sperm from MBS mice tended to have fewer and less extreme morphological abnormalities (Figure 2D). The proportion of intermediate and abnormal nuclei did not increase in either strain with increasing hypoxia exposure duration. We detected no significant differences in the proportion of abnormal (Wilcoxon rank-sum test, p=0.145), intermediate (p=0.112), or normal nuclei (p=0.525) between populations. Overall, this suggests that hypoxia exposure *per se* has a minimal effect on mature sperm morphology across *P. maniculatus.* Additionally, our results suggest that the impaired male reproductive physiology we observed in highlanders results from impairment of male reproductive physiology earlier in spermatogenesis rather than later during the final stages of sperm maturation.

**Figure 2:**
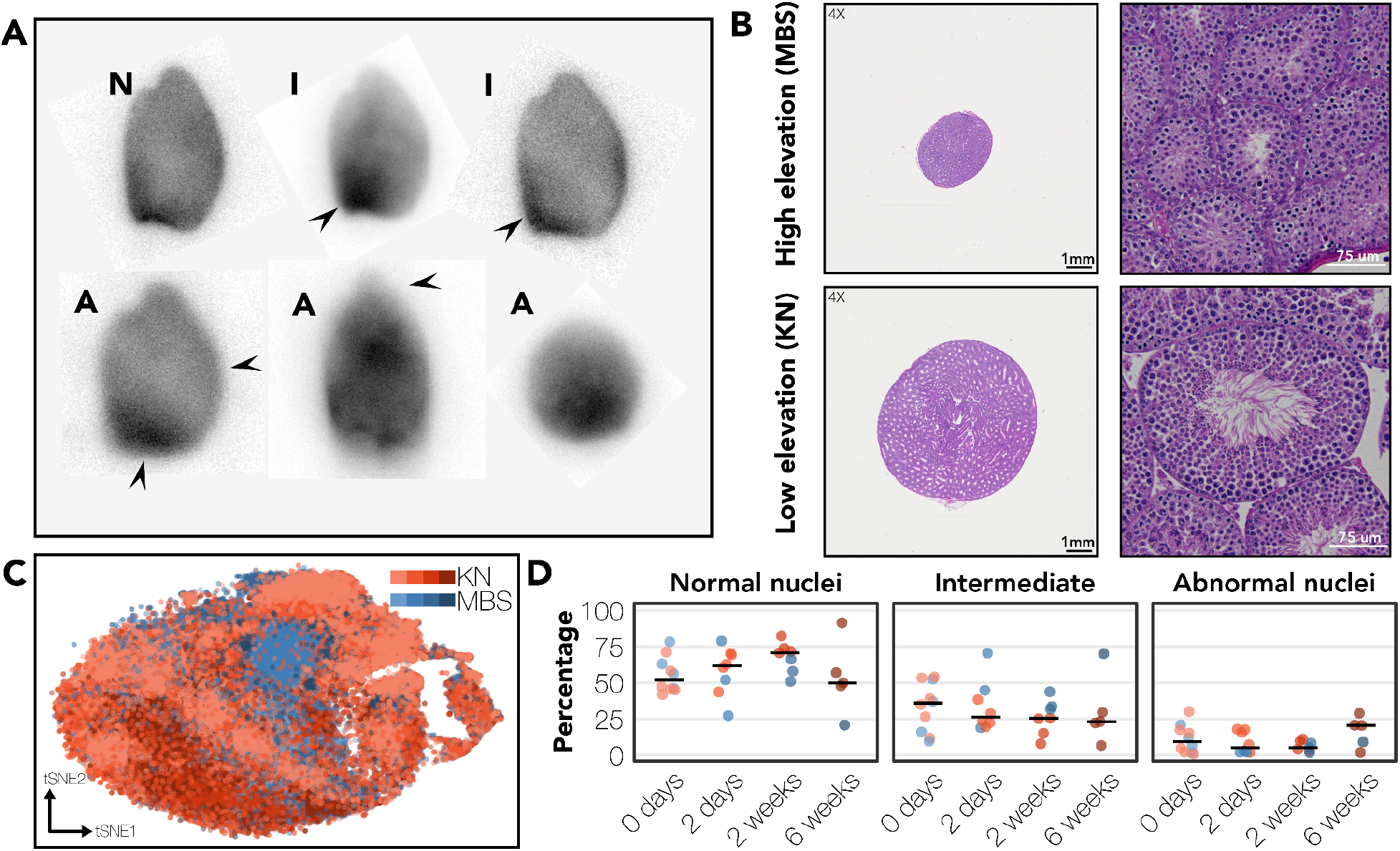
Negligible disruption of spermiogenesis in response to hypoxia despite significant differences in testis architecture between strains in deer mice exposed to hypoxia. (A) Representative DAPI-stained nuclei from sperm morphological groups illustrating normal (N), intermediate (I) and abnormal nuclei. Arrows indicate regions that correlate with spermiogenic perturbation showing increased compression of nucleus and features that correlate with differences in spermiogenic machinery function (Mandawala et al., 2026). (B) H&E stained testes sections from lowland and highland mice at six weeks of hypoxia exposure at 4X (left) and 40x (right). Representative sections of a fertile KN deer mouse and a sterile MBS deer mouse were imaged with a Cytation 5 cell imaging multi-mode reader (BioTek) and stitched 40X images with ImageJ v.1.54p (Preibisch et al., 2009). (C) tSNE analysis of morphological profiles with significant differences between populations ( 24/300 profile measureswere excluded because their Bonferroni-adjusted p-values > 0.05.) colored by hypoxia treatment and strain. Depth of color indicates hypoxic acclimation treatment with darker colors indicating longer duration of hypoxia exposure. Across samples, mature sperm shows broad clustering of morphologies, but a gradient between KN and MBS. (D) The proportion of mature sperm for each individual assigned to each morphological grouping.

### Differential gene expression in testes

To characterize gene expression differences associated with hypoxia exposure in highland and lowland mice, we sequenced testis RNA from deer mice in our focal comparison. Strain of origin was the major factor driving overall gene expression differences across samples (MDS 1: explained 11.1% of variation; Figure 1). By contrast, there was a negligible impact of hypoxia treatment on overall expression differences. Consistent with this, we detected many genes with a strain-specific (MBS vs KN) effect on gene expression (484 genes; 250 upregulated and 234 downregulated in MBS; Table S8). The strain-specific DE genes were enriched for GO terms related to cell signaling receptor activity, localized largely to the cell periphery (Table S9; Figure S4). To prioritize candidate genes related to the evolution of testes phenotypes across strains, we identified whether strain-specific DE genes were previously documented as having higher allele frequency differentiation in MBS relative to lowland conspecifics (*i.e.,* “PBS outliers” from Schweizer et al., 2021; Wilsterman et al., 2023). There were 27 genes that met this criterion (18 upregulated and 9 downregulated). This gene set was not significantly enriched for any GO terms though it does contain many potential gene candidates underlying evolved differences in male fertility, such as those involved in testosterone synthesis (*e.g.,* LOC102913629/*Hsd3b1*) and tissue remodeling (*e.g., Mmp12*).

In contrast to strain-of-origin effects, hypoxia exposure had a minimal effect on overall expression differences within each strain, especially for the lowland strain. For highland mice, overall expression differences were significantly correlated with normalized sperm count (MDS dim 1: ρ = = 0.68; p = 0.002; MDS dim 2: ρ = 0.62; p = 0.008). Accordingly, there was only one gene with either acclimation-specific (2 days: one gene LOC143271886]; 2 weeks: 0 genes; and 6 weeks: 0 genes; Table S8) or strain-by-acclimation effects on differential gene expression (0 DE genes). None of the genes showing hypoxia exposure effects were genome-wide outliers for sequence differentiation between highlanders and lowlanders (Schweizer et al., 2019; Wilsterman et al., 2023). The lack of genes exhibiting strain x hypoxia exposure interaction effects was likely due to both the low variability in overall expression of lowland testes across exposure treatments and the high across- and within-treatment variability in spermatogenic output in highland testes.

### Cell composition drives gene expression differences in high-elevation deer mice testes

Given the correlation between normalized sperm count and overall gene expression that we observed in highland mice, we tested for changes in the cellular composition of testes across highlanders in different hypoxia exposure treatments. We used a deconvolution approach to estimate cell type proportions from our bulk RNAseq data using a house mouse single cell RNAseq testis atlas (Green et al., 2018). This approach revealed shifts in relative cell type proportions in highland but not lowland mice across different hypoxia treatments (Figure 3). We used beta regression to model cell type proportion for four cell types as a function of strain and normalized sperm count. Specifically, we tested for shifts in cell type proportion for a somatic support cell type (Sertoli cells), mitotic precursor cells (spermatogonia), meiotic cells (spermatocytes), and post-meiotic cells (elongating spermatids). We found a significant strain and strain x sperm count interaction for three of the four cell types: Sertoli cells, spermatogonia, and elongating spermatids, but not for spermatocytes (Figure 3). Our results are consistent with shifts in cell type proportions across lowland and highland mice as well as shifts towards fewer late meiotic cell types in the testes in sterile highland individuals. This pattern suggests a dramatic reorganization of testes cell populations in highland mice exposed to hypoxia.

**Figure 3:**
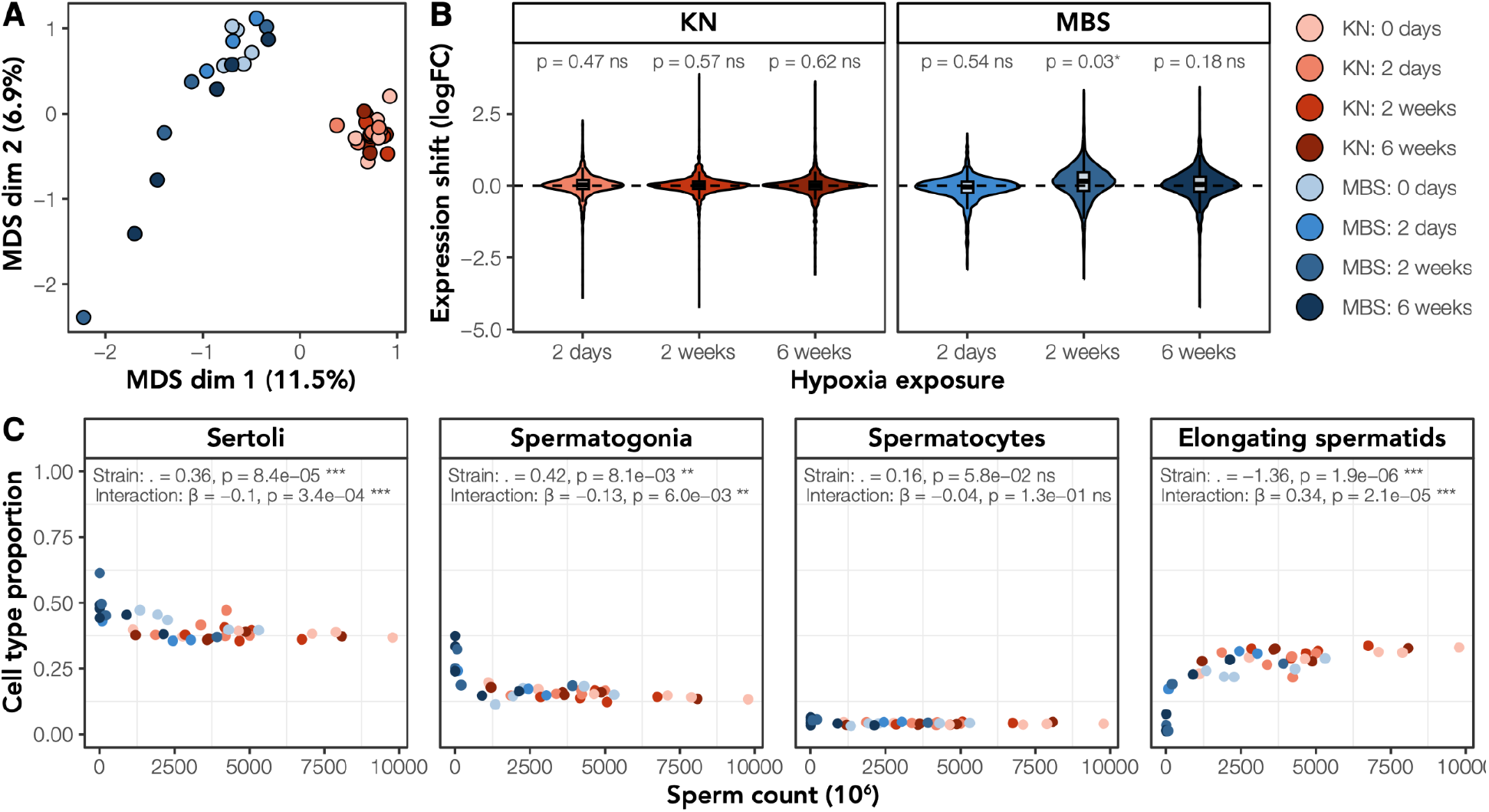
Expression patterns indicate upregulation of apoptosis and shifts in testicular cell composition in highland males exposed to 2 weeks of hypoxia. (A) MDS plots of distances among deer mouse testes samples calculated with the top 1,000 genes that distinguish each sample. Strain indicated by color with Blue = MBS; Red = KN. Depth of color indicates hypoxic acclimation treatment with darker colors indicating longer duration of hypoxia exposure. Axis labels indicate the proportion of variance explained by each logFC dimension. (B) Shifts in expression of apoptosis genes under the parent apoptosis GO term (GO:0006915) in highland but not lowland deer mice with hypoxia exposure. For each hypoxia exposure group, we assessed whether the apoptosis gene set as a whole showed a coordinated shift in expression relative to each strain-specific control using *cameraPR* in limma (Ritchie et al., 2015) and corrected for multiple testing using FDR (Benjamini & Hochberg, 1995) (C) Normalized sperm count and estimated adjusted cell type proportion inferred by deconvolution using MuSiC (X. Wang et al., 2019) for each individual colored by strain and duration of hypoxia exposure for Sertoli cells (support cells), Spermatogonia (pre-meiotic), Spermatocytes (meiotic), elongating spermatids (post-meiotic). Cell type proportions were adjusted with the Smithson-Verkuilen transformation (Smithson & Verkuilen, 2006). Significance results of a beta regression strain, normalized sperm count, and their interaction indicated for each cell type.

To complement this analysis, we also tested for shifts in the overall expression of genes under the GO terms for apoptosis (GO term: 0006915) across hypoxia exposure groups within each strain relative to control mice (0 days exposure). Only highland mice exposed to 2 weeks of hypoxia exhibited a coordinated shift in expression of apoptosis genes with 1022/1632 genes showing upregulation of expression relative to 0 days of hypoxia exposure (mean logFC = 0.11; p-adjusted = 0.03). This peak in expression of apoptosis genes around two weeks could explain the decrease in mature meiotic cell types in the highland mice exposed to hypoxia.

### Gene co-expression networks in the testes and evolution of the hypoxia response pathway

Using WGCNA, we assigned 15,285 testes genes to six co-expression modules (modules “blue”, “brown”, “green”, “red”, “turquoise”, and “yellow”); the remaining 5,286 genes were unassigned (the “grey” module). Modules ranged in size from 347 - 5,658 genes (Figure 4). Because of the high correlation among male reproductive phenotypes (Figure S2), we chose one representative phenotype, normalized sperm count, to identify modules with statistically significant correlations between module eigengene and reproductive physiology. We identified three modules with such correlations [modules “blue” (ρ = -0.49; q = 0.004); “brown” (ρ = -0.70; q < 0.001); and “turquoise” (ρ = 0.61; q < 0.001)] (Figure 4; Table S10). We also used linear mixed effects modeling to test for the effect of hypoxia exposure, strain, or their interaction on module expression. No modules exhibited differentiation between strains, but three modules (“brown”, “turqouise”, and “yellow”) showed nominal strain × treatment effects (p < 0.05, driven mainly by MBS at 2 weeks and 6 weeks) that were not significant after FDR correction (Table S11; Figure 4).

**Figure 4:**
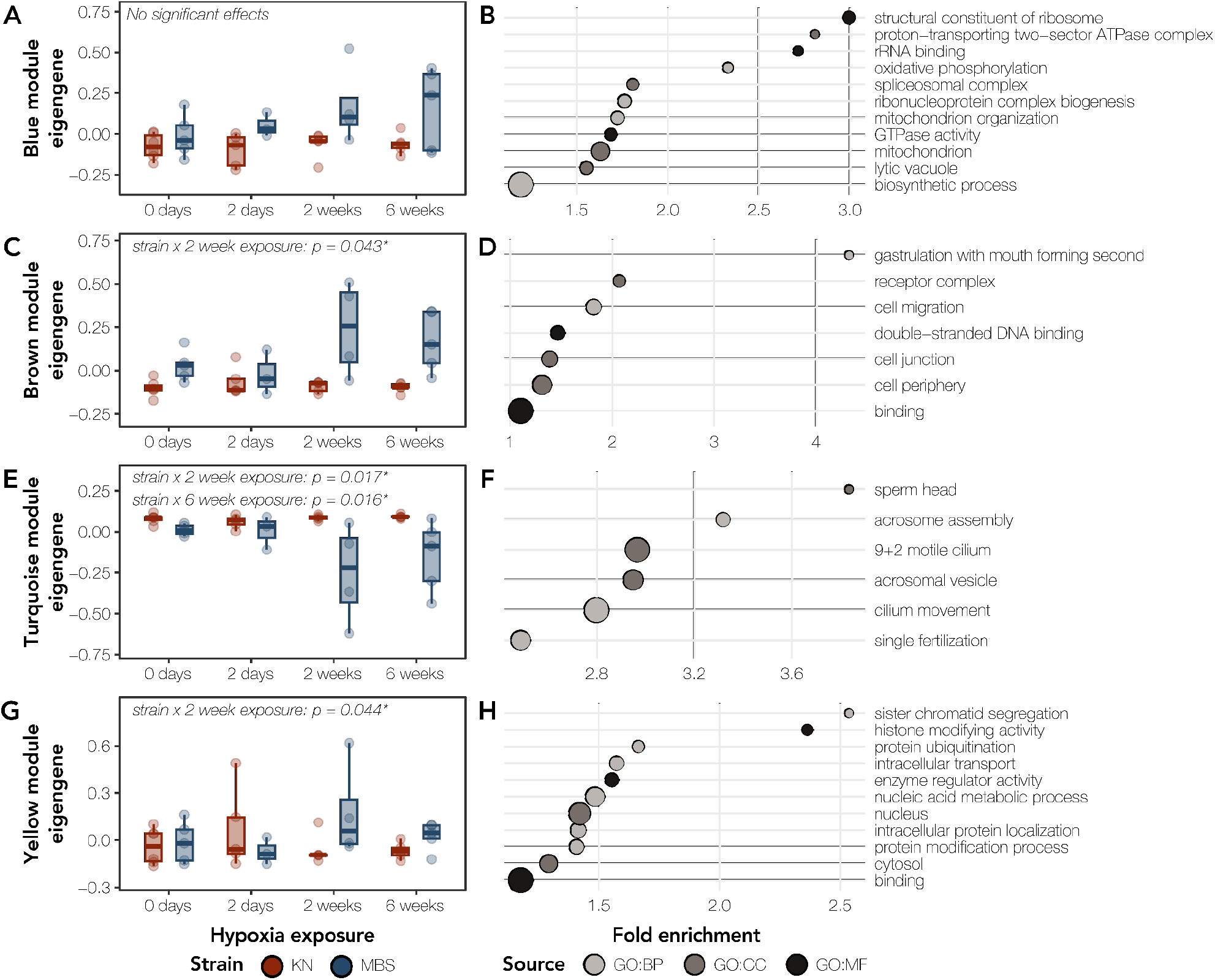
Gene co-expression network modules correlated with sperm count are differentially regulated across hypoxia exposure in highlanders. (A), (C), (E), (G) Module eigengene expression across hypoxia exposure treatments for MBS (blue) and KN (red) deer mice for four testes modules correlated with normalized sperm count or with significant strain x hypoxia exposure effects. Uncorrected p-values correspond to significant effects in linear models of strain, hypoxia exposure, and their interaction (see Methods for more details). (B), (D), (F), and (H) GO terms significantly enriched for genes assigned to each WGCNA module colored by term source (BP: Biological Process, CC: Cellular Component, MF: Molecular Function). Included terms selected by the highlight function within gProfiler2 which collapses GO terms in a two-step clustering algorithm (Kolberg et al., 2020). Point size corresponds to the number of genes belonging to each GO term, and terms are plotted by the fold enrichment of the GO term in the dataset relative to the provided gene backgrounds.

All four testes modules with significant correlations with normalized sperm count and high correlations in our linear mixed effects models were enriched for functional terms based on GO enrichment analysis (Figure 4). Two of these modules were most clearly enriched for GO terms related to spermatogenesis: the “turquoise” and “yellow” modules. The “turquoise” module was enriched for GO terms related to late-stage spermatogenesis which included the terms “sperm head”, “cilium movement”, and “acrosome assembly”, while the “yellow” module was enriched for processes related to sister chromatid segregation, histone modification, and nucleic acid metabolism. For each of these modules, we identified “hub genes” whose expression was most highly correlated with other genes within the same module. We identified highly interconnected hub genes in each module and screened each module for genome-wide outliers for sequence differentiation between highlanders and lowlanders (Schweizer et al., 2019; Wilsterman et al., 2023). While 29/671 of these hub genes had elevated sequence differentiation between highlanders and lowlanders, none were differentially expressed in any contrast in the testes (Table S12).

Finally, we sought to characterize gene expression patterns of key regulators of the hypoxia response pathway across populations and hypoxia exposure treatments. *Epas1* and *Hif1a* were assigned to the same testes module (“brown”) while *Vegfb* was assigned to a different module (“blue”). Of the three genes, only *Hif1a* was in the top 5% of intra-module connectivity in its assigned module (*Epas1 =* 0.80; *Hif1a* = 0.90; *Vegfb =* 0.80). While none of the three genes were differentially expressed in any contrast, highland mice exposed to longer durations of hypoxia (2 weeks and 6 weeks) had elevated expression of all three genes relative to all lowlanders and control highlanders (Figure S5).

## Discussion

Despite its clear importance to fitness, we have a limited understanding of how male reproductive physiology has evolved in response to the reduced oxygen availability at high elevations (but see H.-H. Wang et al., 2026). In this study, we used wild-derived strains of highland and lowland populations of the North American deer mouse, *P. maniculatus,* and a low-elevation strain of the white-footed mouse, *P. leucopus*, to test how male reproductive function has evolved across populations and in response to hypoxic stress. Given the widely-reported negative but primarily transient effects of hypobaric hypoxia (S. Li & Yang, 2022; Liao et al., 2010; Saxena, 1995; Tao et al., 2022; Verratti et al., 2016, 2022), we expected high-elevation populations of deer mice to be buffered from the detrimental impacts of hypoxia exposure on male reproduction. Contrary to this expectation, highland deer mice showed both overall decreased fertility relative to low-elevation counterparts and an increased sensitivity to hypoxic stress, including complete sterility in as little as two weeks under hypoxia. This unexpected result suggests that adaptation to chronic hypoxia incurs a reproductive cost in highland deer mice. Below, we explore possible evolutionary explanations for this counterintuitive physiological response.

### Differences in male fertility between high- and low-elevation populations

Deer mice are an exemplar species of a promiscuous mating system with documented high levels of sperm competition (Fisher et al., 2014; Gozashti et al., 2021; McLean et al., 2022; Voss & Nachman, 2025; Weber & Fisher, 2023), although we note that most studies have focused on a single lab-derived strain (BW), and variation across the *P. maniculatus* range remains to be quantified. Because of the documented effects of sperm competition in other *P. maniculatus* populations, the reduced baseline male fertility in both independently-derived high-elevation strains is striking and suggests that reduced baseline fertility is a general characteristic of high-elevation populations rather than a unique feature of our MBS strain. It is possible that the lower baseline fertility of high-elevation deer mice results from the use of inbred colonies. However, both of the low-elevation stock center lines (LL and BW) have been maintained in captivity for as many or more generations than our focal high-elevation strain (Lucius et al., 2021), suggesting that colony effects alone cannot fully explain lower baseline fertility of highlanders. Our male fertility data from the CSU colony further confirmed this reduced baseline fertility, but suggested that baseline fertility is flexible in response to social environment (male versus female exposure). Importantly, males from both lowland and highland strains had similar times to first litter and similar litter sizes (KW, data not shown). Together, our findings suggest that highlander deer mice have reduced baseline fertility but may maintain an equal capacity for reproductive competence in the absence of hypoxic exposure.

Two possible mechanisms could result in the apparent decreased fertility of high-elevation deer mice: relaxation of selection on male reproductive output and environmental and ecological differences between laboratory and high-elevation environments. Relative testes weight is reduced in both high-elevation strains relative to the low-elevation strains, which is consistent with a relaxation of sperm competition in highlanders (Parker, 2016; Ramm et al., 2014). While deer mice are generally assumed to have a promiscuous mating system (Fisher et al., 2014; McLean et al., 2022; Voss & Nachman, 2025; Weber & Fisher, 2023), there is little known about the mating system and social relationships in the wild for high-elevation species (McLean et al., 2022; Weiner et al., 1984), although at least three primarily low-elevation species in the genus have evolved a monogamous mating system (*P. californicus*, *P. eremicus*, and *P. polionotus*; Gozashti et al., 2021; Niepoth et al., 2024; Voss & Nachman, 2025).

One important caveat to this possibility is that highland deer mice had less variable sperm morphology than lowlanders, and typically, reduced sperm competition leads to more variable sperm (Breed et al., 2007; Calhim et al., 2007; Immler et al., 2010; Rowe et al., 2015; Suttle et al., 1988; Thitipramote et al., 2011). The subtle morphological differences in overall nuclear shape between our two deer mouse strains resemble other rodent populations including *Mus musculus musculus* and the less inbred strains of *Mus musculus domesticus* (Skinner et al., 2019). The normal, intermediate, and abnormal sperm nuclear morphology categories are very similar in feature to the same categories in (Mandawala et al., 2026) showing similar signs of spermiogenic perturbation, despite the differing fundamental morphology of the falciform deer mice and the spatulate pig sperm. The resultant phenotypes differ due to the aforementioned falciform-spatulate difference but share similar evidence for differences in compression and acrosome-acroplaxome-manchette complex formation during spermiogenesis. The signs of increased nuclear condensation in regions compressed during spermiogenesis support the idea of a similar developmental mechanism (Figure 2A). Because our highland deer mice had less variable sperm morphology than lowlanders, our results are consistent with selective constraint on sperm function and not relaxed selection. However, *in vivo* monitoring of burrows and paternity tracking are needed to address the likelihood of a shift to higher rates of monogamy.

A second possible explanation for lower baseline fertility of highland mice is that the reduced baseline fertility we observed in our lab populations is not reflective of baseline fertility in nature during peak breeding season. Our study replicated only one dimension of the complex environments of wild high-elevation deer mice, which would also experience colder, longer winters (McLean & Guralnick, 2021), lower primary productivity (Dillon & Conway, 2021), among other effects like heavy metal exposure (Clements et al., 2000). Prior work has demonstrated decreased male fertility in *Peromyscus* species exposed to reduced photoperiod and restricted water and green food availability (Nelson, 1993; Nelson et al., 1995, 1997). Further, we intentionally socially isolated males several weeks before the focal experiment to mitigate dominance effects (Snyder, 1967). The mice in our experiment were also born and raised to adulthood at a lower elevation than their native range. It is possible that the reduced fertility of high-elevation deer mice is a transient phenotype that is developmentally (dependent on oxygen availability during development), temporally (across the breeding season), and socially (in the presence of mates and competitors) plastic. In support of this hypothesis, our results demonstrate that males exposed to a female have larger testes than those housed with same-sex siblings (Figure 1F). Therefore, baseline fertility of highland males may be elevated in the wild during breeding season, when resources are abundant and competition to secure paternity is highest. This plasticity in reproductive phenotype would be advantageous in times where maintaining spermatogenesis is costly, and the probability of paternity is low. In sum, evidence of large effective population size in highland mice (Schweizer et al., 2021) and data reporting similar time to first litter and litter size suggest that the reduction in baseline fertility is either unlikely to detrimentally impact population size or is not reflective of fertility during the peak breeding season.

### Sensitivity of high-elevation male reproductive physiology to hypobaric hypoxic stress

Our second main finding was that hypoxia exposure negatively impacted male fertility in high-elevation mice. We expected to observe negative effects of hypoxic stress on male reproductive physiology in our lowland deer mice, consistent with previous work in lab models (S. Li & Yang, 2022; Liao et al., 2010; Saxena, 1995; Tao et al., 2022). Instead, we found no significant effects of hypoxia exposure on relative testes weight, seminal vesicle weight, and average seminiferous tubule area. Similar to a previous study of hypobaric hypoxia challenges in house mice, we even observed a trend towards increased relative testes weight in our KN strain across hypoxia exposure treatments (S. Li & Yang, 2022), hypothesized to be consistent with a swelling of the testis interstitium in response to hypoxic challenge. The lack of impairment on lowland reproductive physiology in response to hypoxic exposure could be related to our study elevation (∼1,640 meters) and any impacts on male reproductive phenotypes that occurred prior to the beginning of the experimental trials. Fine scale studies into which simulated partial pressures of oxygen result in impaired male reproductive physiology would help clarify this possibility. Finally, there may have been additional fine-grained negative effects on spermatogenesis in low-elevation mice that could not be observed in this study.

In stark contrast, both histological changes in testes architecture and our gene expression results are consistent with a regressed testicular phenotype in sterile highland males, as occurs in seasonally breeding mammals (Beltrán-Frutos et al., 2022; Brown, 1966; Korytko et al., 1995; Weiner et al., 1984). Relative testes weight, normalized sperm counts, and seminiferous tubule area all decreased in response to hypoxia in highlanders. Deconvolution of testes bulk RNAseq data suggested a restructuring of testicular cell types including decreases in late meiotic cell types like spermatids beginning at two weeks of hypoxia exposure with corresponding increases in spermatogonia and support cells like Sertoli cells. Further, the increase in expression of apoptosis genes at two weeks of hypoxia exposure is consistent with programmed restructuring of the testes and active clearing of mature spermatogenic cell types, which likely resulted in the observed azoospermia. Our expression results are consistent with disruption at or before the completion of meiosis which resembles increases in apoptosis during seasonal testicular regression in deer mice (Young et al., 1999, 2001) and other mammals (An et al., 2025; Bueno et al., 2024). Studies from house mice suggest that recovery from hypobaric hypoxia exposure can take at least six to nine weeks (Bai et al., 2018; S. Li & Yang, 2022; Z. Li et al., 2021). Future experiments with extended hypoxia exposure, comparisons to wild high-elevation mice in the non-breeding season, and experimental trials where individuals are returned to normobaria following hypoxia exposure would provide further evidence that the hypoxia exposure effect on male reproductive physiology we observed is plastic and consistent with testicular regression. Whether the regenerative capacity of spermatogenic stem cells is altered in high-elevation deer mice is another open question these future studies would help address.

We hypothesize that the increased sensitivity to hypoxia exposure in high-elevation deer mice reflects an adaptive reproductive strategy to triage energy expenditure under stressful conditions. Extreme environments, including high elevation, can force trade-offs between survival and reproduction (Hubert et al., 2026; Smith & Charnov, 2001; Yahr & Kessler, 1975). As a result, many mammals, including *Peromyscus* (Nelson, 1993; Nelson et al., 1995; Young et al., 2000), have evolved temporally plastic reproductive strategies including seasonal shutdowns and restarts of both spermatogenesis (Beltrán-Frutos et al., 2022; Jiménez et al., 2015) and the estrous cycle (Bronson & Heideman, 1994; Chemineau et al., 2008). Increased sensitivity to environmental stressors would permit high-elevation males to invest in immediate aerobic metabolic needs over reproduction. At such extreme elevations, foraging deer mice live in near constant risk of hypothermia even in summer, placing severe pressure on the ability to perform aerobic metabolic activities such as thermoregulation (Hayes and O’Connor 1999). Accordingly, highland deer mice have strongly elevated thermogenic abilities relative to lowlanders (Cheviron et al. 2012; Lui et al. 2015; Velotta et al. 2016; Tate et al. 2020;), which is under intense selection at high elevation in some years (Hayes and O’Connor 1999). The extreme energetic demand of high elevation is also known to place limits on energetic tasks related to the development of the thermoregulatory system (Robertson and McLelland; Velotta et al. 2020). Intriguingly, the complete azoospermia we observed in some highlanders under hypoxia exposure was variable across study individuals. If there is standing variation in reproductive plasticity across highland individuals and if this variation is heritable, it may provide the substrate for selection on male reproductive plasticity.

Future experiments exposing mice to alternative chronic stressors (*e.g.,* prolonged cold, predatory threat) would help to clarify whether our results can be explained by stress in general or hypoxia specifically. It seems unlikely that testicular regression should be sensitive soley to hypoxia, given that limited oxygen availability is invariably extreme at high elevations. Thus, we feel our results have uncovered a universal response to stress triggered by the rapid switch from normobaria to hypobaria and not necessarily from the lack of oxygen by itself.

### The regulatory underpinnings of hypoxia-induced sterility in highland deer mice

Both the hypoxia response cascade and germ cell differentiation share an overlapping gene set and specific molecular pathways including HIF and VEGF signaling, mTOR metabolism, and oxidative stress signaling (Lord, 2025). *Hif1a*, *Epas1*, and *Vegfb* are important genes in the hypoxia response cascade, but we did not observe differential expression of these candidate genes in response to strain, hypoxia exposure duration, or their interaction. Recent evidence from house mice suggests that *Epas1* expression is critical for spermatogenic stem cell renewal (Bernstein et al., 2023) which would likely play a critical role in recrudescence following testicular regression. Although these genes have documented roles in other physiological adaptations to high elevation (Schweizer et al., 2019), whether they exhibit differential regulation or pleiotropic effects of coding or regulatory mutations in specific testes cell populations remains an open question.

In contrast to known regulators of the hypoxia response cascade, our differential expression analysis and gene co-expression networks point to candidate genes that may underlie the shifts in reproductive physiology that we have documented here. These include two genes with high sequence differentiation between highlanders and lowlanders and show the largest logFC between strains in the testes: *Hsd3b1* and *MMP12*. *Hsd3b1* encodes the enzyme 3β-hydroxysteroid dehydrogenase (Matsuyama & DeFalco, 2023), which is part of the testosterone synthesis pathway in Leydig cells (Xue et al., 2022), and transgenic knockouts in zebrafish result in extensive endocrine and reproductive disruption (L. He et al., 2026). *MMP12* (matrix metalloproteinase-12) is expressed in Sertoli cells (C. Chen et al., 2005) and may help with stem cell recruitment and migration (Hess et al., 2006). Despite the presence of promising candidates, we note that many hub genes in hypoxia-responsive or trait-correlated modules of interest were not differentially expressed, and many of the largest effect DE genes (as measured by logFC) were not genome-wide outliers for sequence differentiation between highlanders and lowlanders. Our study relied on bulk RNAseq data which is sensitive to shifts in cellular composition among treatments (Hunnicutt et al., 2022) like those we documented with our cell composition analysis. The cell composition differences between the fertile and non-fertile mice in hypoxia exposure treatments likely increased expression variance within treatment and reduced our power to estimate differential expression. We present these candidate genes with the caveat that future experiments using single cell RNAseq data from these testes and genetic mapping crosses would allow finer scale resolution into the molecular basis of these reproductive phenotypes.

## Conclusions

High-elevation environments impose challenges to the integrated physiological systems that must work together to orchestrate organismal function. Here, we document that adaptation to high elevation in *P. maniculatus* does not result in buffered male reproductive physiology to hypoxia exposure, and that plastic impairment of male reproductive physiology may serve as a solution to the opposing energetic demands of survival and thermogenesis with reproductive output. We still have much to learn about how organism function evolves across multivariate trait space given constraints poised by other physiological processes, and it remains unclear how other environmental, ecological, and social factors might interact with hypoxic conditions in the wild to shape male fertility. Our study underscores the importance of accounting for the full environmental complexity of extreme environments and of coordinated physiological responses to further our understanding of adaptation.

## Data availability

All code used in the generation of figures in this manuscript is available at https://github.com/KelsieHunnicutt/Hunnicutt_et_al_2026_Deer_mouse_male_fertility_across_el <u>evation</u>. The sequencing data generated in this study have been deposited in the NCBI BioProject database under accession number PRJNA1544162.

## Acknowledgements

We would like to thank members of the Velotta Lab for feedback on this project, and Ivan Kovanda for support using the University of Denver Research Data Analysis Cluster. The content of this manuscript is solely the responsibility of the authors and does not necessarily represent the official views of the National Institutes of Health or the National Science Foundation. We thank Zac Cheviron for providing funding and animal care and husbandry for the original founders for MBS and KN animals used in the acclimation experiments.

## Funding

KEH was partially supported by a National Science Foundation Graduate Research Fellowship (DGE-2034612) and NIH NIGMS (F32GM163371) throughout the duration of this project. Funding for JPV, KEH, ESG, MQT, and LKC provided by an NIH NIGMS (R35GM147459) grant to JPV. ELL was supported by a National Science Foundation grant to ELL (DEB-2337629). Funding for KW and MYJ was provided by NIH NICHD (R01 HD114615), NSF IOS (2512875), and a Boettcher Foundation Webb-Waring Biomedical Research Award.

## Author Contributions

KEH and JPV conceived of and designed the study. KEH, ESG, JP, MT, LC, MYJ, and KW collected the data, and KEH and JCP analyzed the data. KEH wrote and revised the initial manuscript with feedback from ELL and JPV. All authors read and approved the final version of the manuscript.

## Notes

### Competing Interest Statement

The authors have declared no competing interest.

