## Supplemental Figures for "Reduced baseline male fertility and increased sensitivity to hypoxic stress in high-elevation deer mice"

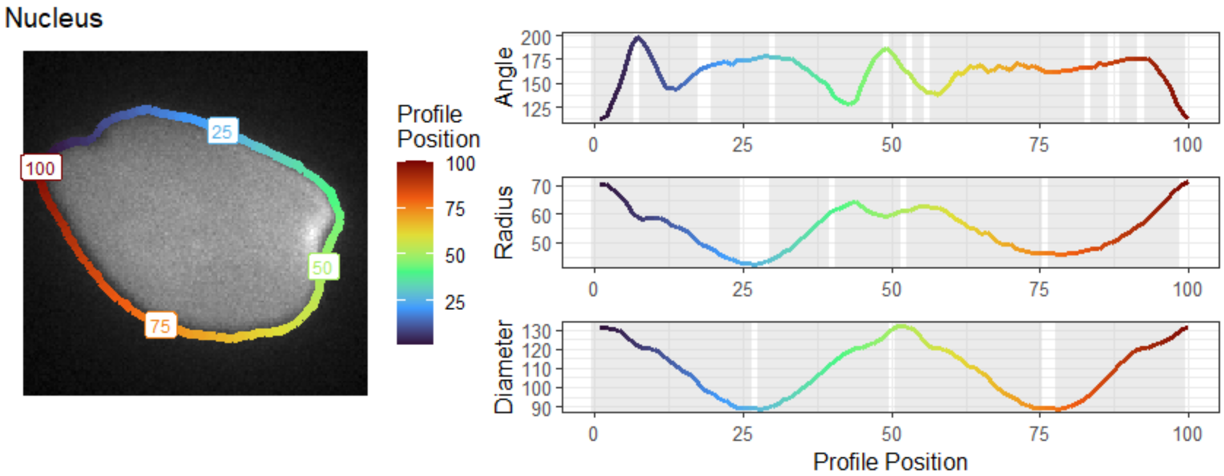


**Figure S1:** Example morphometric profiles with the outline overlaid on an image of a sperm nucleus. Color indicates corresponding profile positions and landmarks. Angle is in degrees, while diameter and radius are in pixels. Significantly different regions between populations have a grey background (P<0.05 with Bonferroni correction).

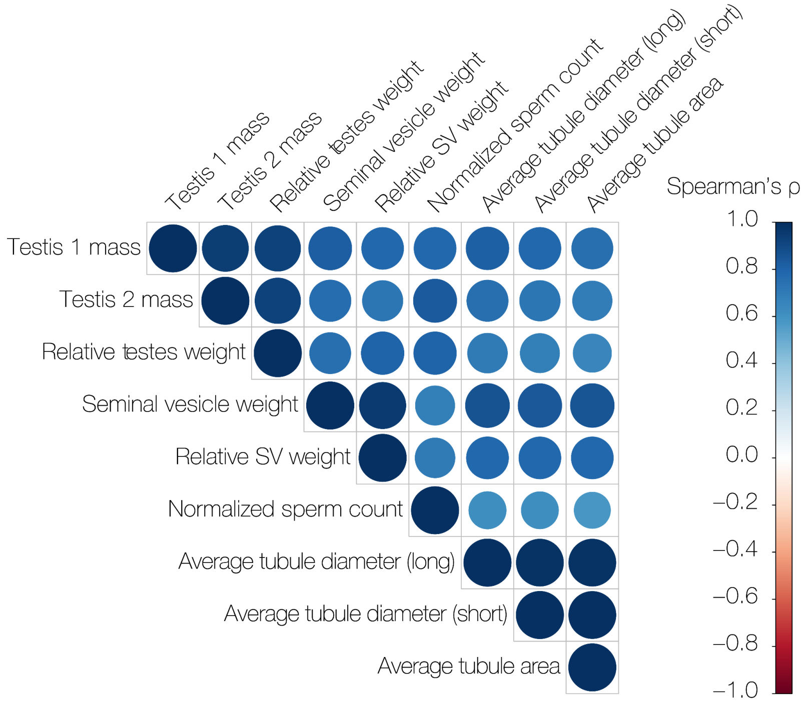


**Figure S2. High positive correlation across male reproductive traits in *P. maniculatus*.** We calculated Spearman's correlations across traits with the *rcorr* function within the *Hmisc* package in R and visualized the correlation matrix with the *corrplot* v0.95 package. Traits included are individual testis weights (mg), relative testis weight (mg/g), seminal vesicle weights (mg), relative seminal vesicle weights (mg/g), normalized sperm count (in million sperm/mL), average seminiferous tubule diameter across short and long axes (um), and average seminiferous tubule area ($mm^2$).

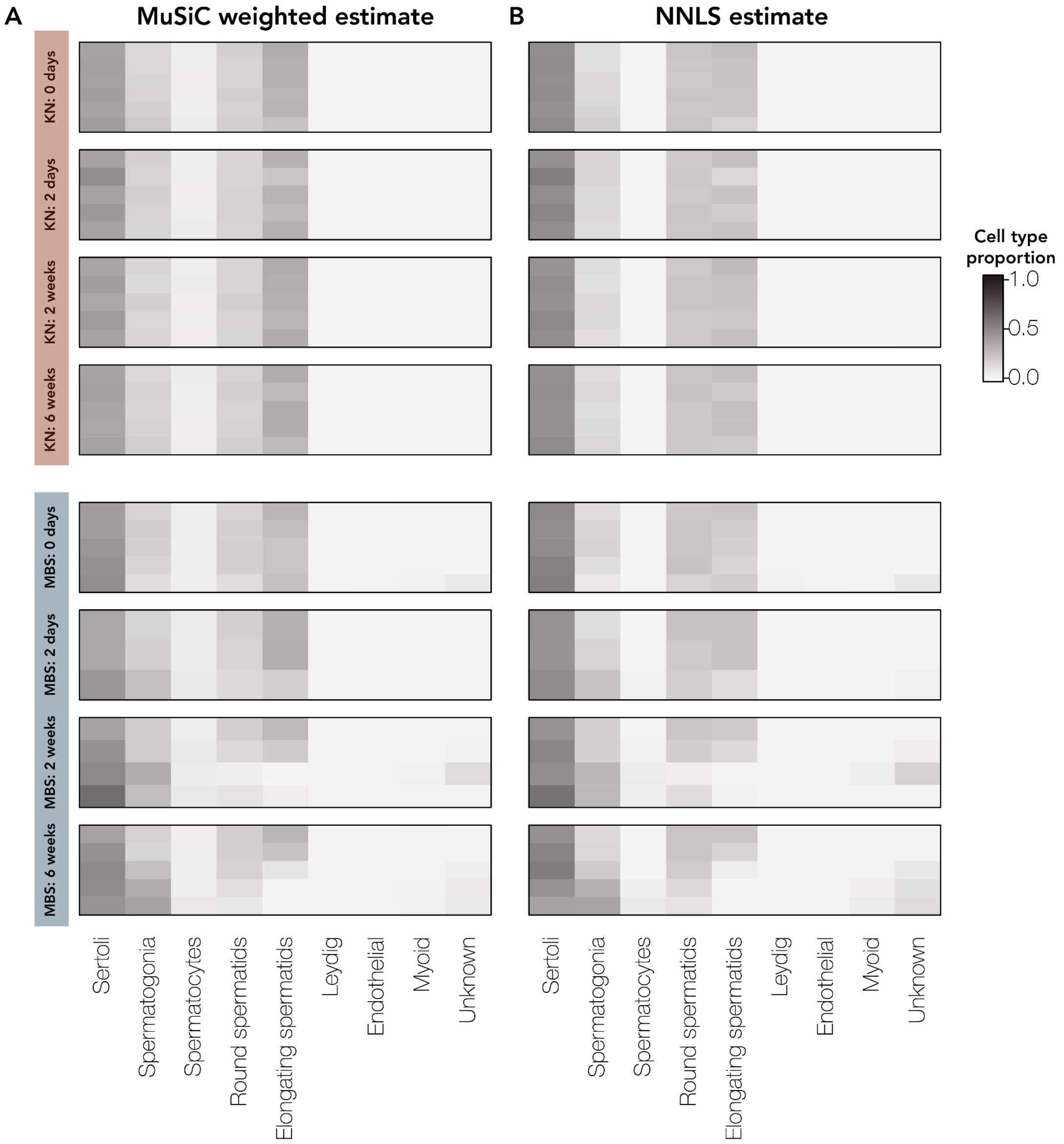


**Figure S3: Cell type proportion estimates across test bulk RNAseq samples.** We estimated cell type proportions for each sample with music_prop in MuSiC (X. Wang et al., 2019) using a house mouse testes single cell atlas (Green et al., 2018) resulting in a MuSiC-weighted estimate (Est.prop.weighted) and a non-negative least squares (NNLS) estimate. Assessed cell types included Sertoli cells, spermatogonia, spermatocytes, round spermatids, and elongating spermatids, Leydig cells, innate lymphocytes, endothelial cells, myoid cells, and an unresolved ("unknown") population. Samples arranged by strain x hypoxia treatment first, then by decreasing normalized sperm count within each box.

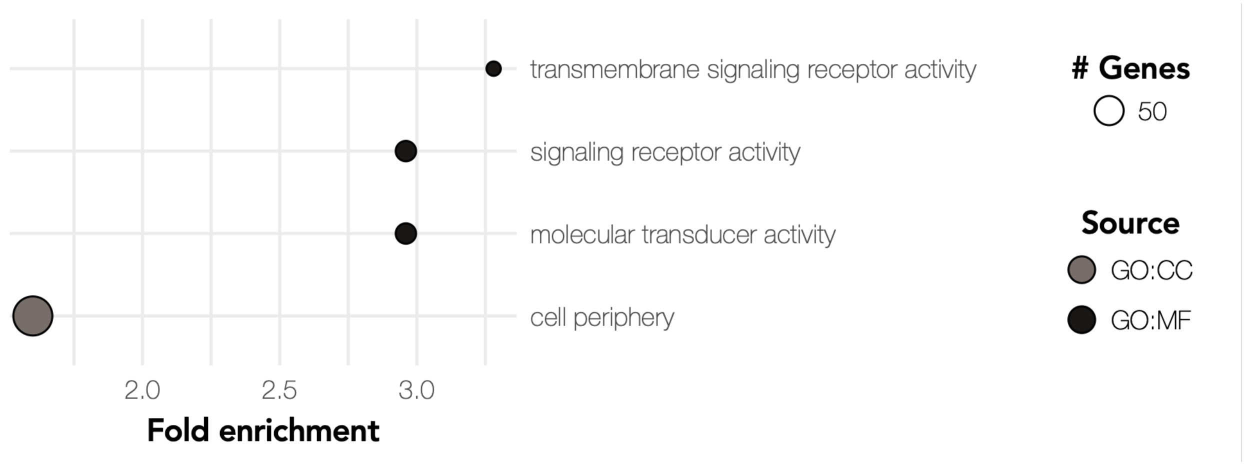


**Figure S4. GO term enrichment of differentially expressed genes between strains.** GO terms significantly enriched for genes with a significant strain effect on differential expression colored by term source (BP: Biological Process, CC: Cellular Component). Point size corresponds to the number of genes belonging to each GO term, and terms are plotted by the fold enrichment of the GO term in the dataset relative to the provided gene backgrounds.

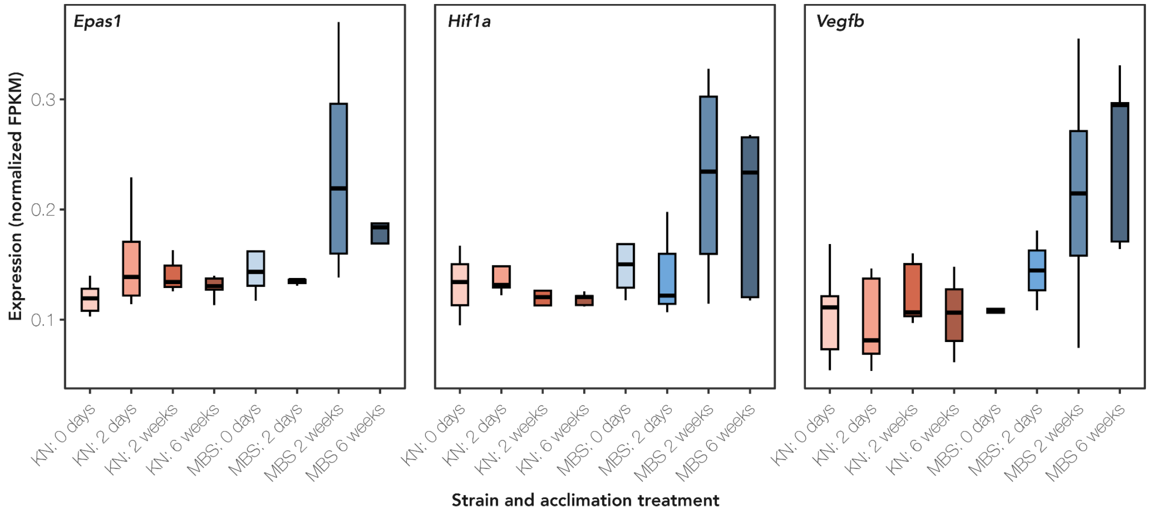


**Figure S5: Expression of key genes in the hypoxia response cascade. (**A) *Epas1*, (B) *Hif1a*, and (C) *Vegfb* expression across strains and hypoxia exposure treatments where expression is a normalized FPKM where the sum of squares equals 1 using the R package vegan (Oksanen et al. 2007). Strain indicated by color with Blue = MBS; Red = KN. Depth of color indicates hypoxic acclimation treatment with darker colors indicating longer duration of hypoxia exposure.
